# Darwinian dynamics over recurrent neural computations for combinatorial problem solving

**DOI:** 10.1101/2020.11.06.372284

**Authors:** Dániel Czégel, Hamza Giaffar, Márton Csillag, Bálint Futó, Eörs Szathmáry

## Abstract

Efficient search in enormous combinatorial spaces is an essential component of intelligence. Humans, for instance, are often found searching for optimal action sequences, linguistic structures and causal explanations. Is there any computational domain that provides good-enough and fast-enough solutions to such a diverse set of problems, yet can be robustly implemented over neural substrates? Based on previous accounts, we propose that a Darwinian process, operating over sequential cycles of imperfect copying and selection of informational patterns, is a promising candidate. It is, in effect, a stochastic parallel search that i) does not need local gradient-like information and ii) redistributes its computational resources from globally bad to globally good solution candidates automatically. Here we demonstrate these concepts in a proof-of-principle model based on dynamical output states of reservoir computers as units of evolution. We show that a population of reservoir computing units, arranged in one or two-dimensional topologies, is capable of maintaining and continually improving upon existing solutions over rugged combinatorial reward landscapes. We also provide a detailed analysis of how neural quantities, such as noise and topology, translate to evolutionary ones, such as mutation rate and population structure. We demonstrate the existence of a sharp error threshold, a neural noise level beyond which information accumulated by an evolutionary process cannot be maintained. We point at the importance of neural representation, akin to genotype-phenotype maps, in determining the efficiency of any evolutionary search in the brain. Novel analysis methods are developed, including neural firing pattern phylogenies that display the unfolding of the process.

## 1 Introduction

Life, with its “endless forms most beautiful” [1], is a result of a Darwinian evolutionary process operating over the enormous representational capacity of chemistry. Any Darwinian process is built on the principles of replicating units, hereditary variation, and selection [2]; beyond the ability to sustain these, a Darwinian process makes no further demands of the underlying substrate. Replication, hereditary variation and selection collectively operate at other levels of biological organization, for example in the mammalian immune system [3], and may well feed on other-than-chemical substrates. The brain’s neural networks are another example of a system that produces seemingly endless beautiful forms, from neural activity to the mental states and actions that they support. The endeavor of *Darwinian neurodynamics* explores the possible ways in which i) Darwinian dynamics might emerge as an effective high-level algorithmic mechanism from plasticity and activity dynamics of neural populations [4, 5] (*how does it work?*) and ii) this high level algorithmic mechanism fits into cognition (*what is it good for?*). This endeavour can be seen as part of a more general pursuit of a theory of high-dimensional adaptations that unifies our understanding of evolutionary and learning processes (see Table 1).

**Table 1:**
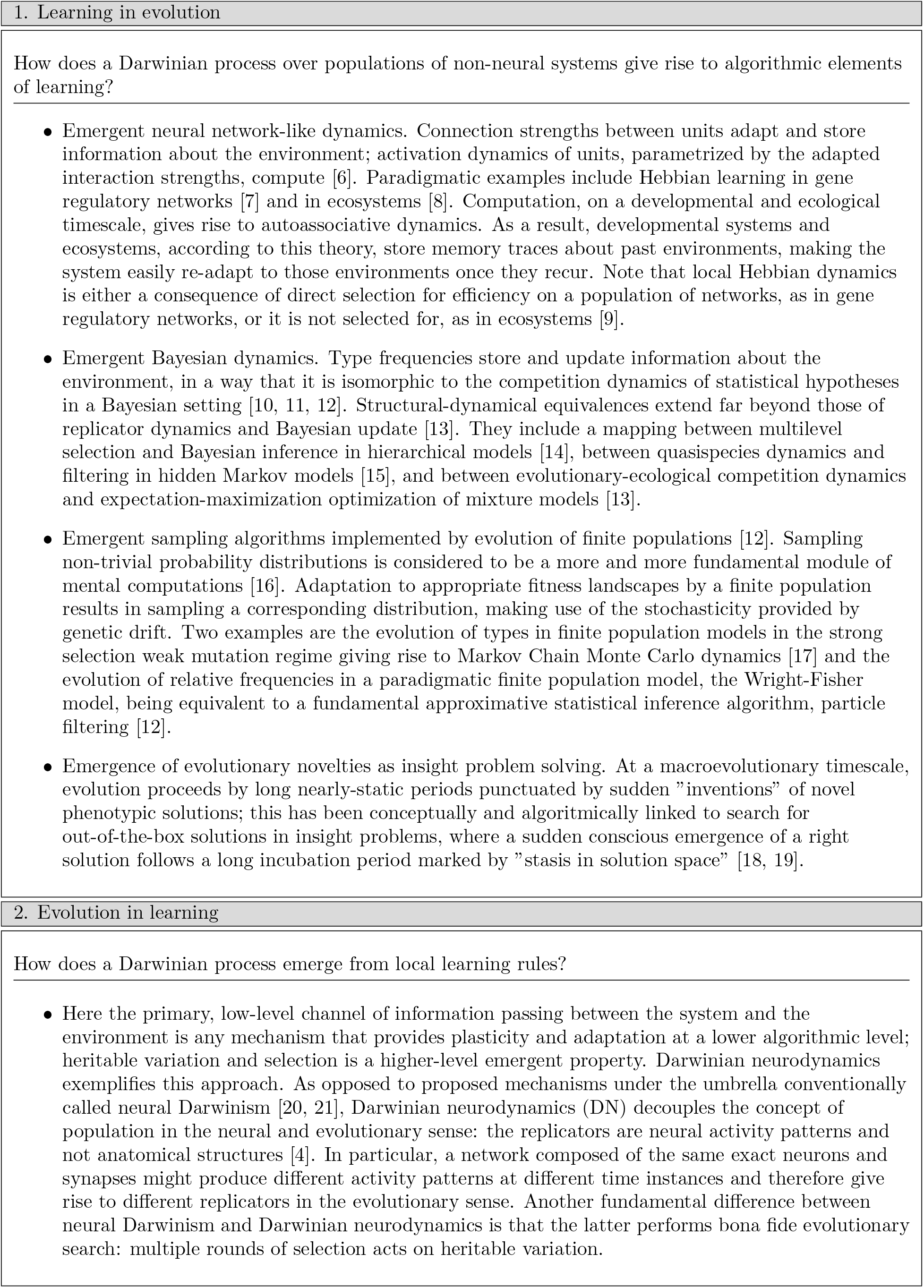
A unified view of evolutionary and learning processes, as part of a theory of high-dimensional adaptations.

In any model of Darwinian neurodynamics (DN), the modeller is required to specify how the ingredients of Darwinian evolution, namely replication, selection, and heritable variation of firing patterns, emerge from lower-level neural activity and synaptic plasticity rules. A related early idea, termed *neural Darwinism*, together with its more recent implementations, considers selective amplification of a pre-existing but possibly adaptive repertoire of computational units [22, 23, 21, 24, 25]. It is halfway between traditional neural adaptation mechanisms, implementing effective high-dimensional hill-climbing in representation space, and a bona fide evolutionary search, with multiple rounds of selection over heritable variation generated by high-fidelity replication [26, 4]. Building on these selectionist ideas, the *neuronal replicator hypothesis* [27] explores possible ways to add the crucial missing ingredient, replication, using exclusively local neural plasticity and activity rules. Neural replicators proposed to date come in two flavours: combinatorial replicators, copying a sequence of low-informational states of neural activity (represented by e.g. bistable neurons), akin to template replication [28, 27], and holistic replicators [29], copying one high-informational state, such as memory traces of autoassociative attractor networks [5]. Three broad categories of *neuronal* units of evolution have been suggested: i) neural activity patterns [5, 28, 27], ii) connectivity, and iii) evolvable neuronal paths [24]. If neuronal replicators exist, their appearance marks a new way information is represented in evolutionary units and transmitted between them - a filial major transition in evolution, as proposed in [30].

DN is similar to other approaches to understanding the computational functions of the mind and its neural correlates in the brain: efficient theoretical frameworks of adaptation, notably reinforcement learning and statistical learning, have been successfully guiding experimentation in neuroscience in recent years. As in the aforementioned theories, in DN, the neural correlates of crucial algorithmic elements, replication, selection and heritable variation, are a priori unknown, emphasizing the importance of hypothesis-driven search. When searching for these algorithmic ingredients of an emergent Darwinian process, three important conceptual issues arise recurrently.

1. **Unlimited hereditary potential**. Long term evolution rests on the ability of novel solutions, generated via mutation, to replicate. If the set of solutions that can be copied with high-enough fidelity is limited, no efficient evolutionary search can take place. This requirement for high enough fidelity copying of a very large range of potential solutions is one that direct in silico models of evolution rarely have to consider, as replication can be achieved simply with arbitrary precision. When the replication mechanism is itself explicitly modelled, the question of its capacity to replicate solutions must be addressed.
2. **Maintenance of information**. Although the appearance of novel beneficial mutations is necessary, it is not sufficient for continuous evolutionary search; information must also be maintained. In particular, the information that is lost through deleterious mutation must (at least) be compensated for by selection. This sets an upper limit on the mutation rate, depending on the selective advantage of the current solution over the surrounding ones.
3. **Representation of phenotype/solution**. That representation really matters is a conclusion arrived at by both evolutionary and learning theory largely independently. In evolutionary theory, phenotypes over which selection acts, are represented by genotypes that replicate. In neurobiology, computational states (e.g. solutions to a given problem) are represented by neural activity patterns. These mappings between genotype/neural activity and phenotype/solution might be highly complex. Representation is important for at least two different reasons. For one, it provides enormous adaptive potential: appropriate transformation of rich-enough representations might render simple algorithms surprisingly powerful. In evolution, in particular, this permits genotypic mutations to be channeled along some, potentially fit, phenotypic directions; this is one of the mechanisms upon which evolvability rests. For the other, we must separate the effect of representation and the algorithm operating on top of it; in particular, the evaluation of the efficiency of algorithms based on a few hand-picked and hand-fabricated representations of a given problem. This issue is unavoidable in DN, as we have at this point no detailed knowledge of how particular mental states are represented in human and animal brains. That is, our map between relevant variables at the computational/behavioral, algorithmic and implementational level remains far from complete.

In this paper, we propose a model of Darwinian neurodynamics based on the emergent replication of *dynamic* patterns of neural activity. Building on the versatility and efficiency of the reservoir computing paradigm, we demonstrate that dynamic activity patterns can be effectively replicated via *training one recurrent network by another*. The two main components of our model that map both to building blocks of evolutionary dynamics and proposed mechanisms of mental search (e.g. in the combinatorial space of action sequences) are *evaluation* of proposed activity patterns and nearly perfect *copying* of them. The latter proceeds via a *teacher* network, i.e. the network that generated the fitter signal, training a *learner* network, i.e. associated to the less fit signal. We show that an ensemble of such recurrent networks, capable of training one another in parallel based on previous evaluation of their output signals, can give rise of an effective evolutionary search that *i*) maintains information about already discovered solutions and *ii*) improves on them occasionally through the appearance and spread of beneficial mutations.

By continually selecting and imperfectly replicating solutions that are better fit to the demands of the cognitive task at hand, and discarding those which don’t meet the muster, DN can be seen as a process of stochastic parallel search with redistribution of resources. In the biosphere, this process of Natural Selection successfully operates in an enormously high dimensional (genomes range in size from ~ 2kb to 10^8^kb), combinatorial (each position can be one of four bases) and compositional (genomes are semantically and syntactically complex) space of genotypes. The solution spaces of problems in many cognitive domains share these characteristics to differing extents. Language is one such domain; indeed an evolutionary computational framework has previously been reported addressing aspects of this faculty [31], in which linguistic constructs (as units of evolution) can be coherently multiplied with hereditary variation under selection to achieve e.g. mutual comprehension in a population. Causal reasoning and action planning are two other examples of cognitive domains characterized by complex search spaces, where it is unclear if local gradient information is useful. To model these complex search spaces in the most generic manner possible, we focus on the capacity of neural-evolutionary dynamics to solve two classes of combinatorial search problems: the Travelling Salesman Problem (TSP) and the tuneably rugged NK landscapes [32]. We demonstrate novel analysis methods that combine approaches from computational neuroscience and phylogenetics, and are generally applicable to any proposed DN architecture.

This endeavour of understanding the implementational, algorithmic and computational features of Darwinian neurodynamics is worthwhile for at least two reasons: i) for its potential as a useful, though not necessarily ubiquitous, computational module in wet brains; a capacity, which, if it exists, has been selected for by biological natural selection, and ii) for its potential as a computational module that is orthogonal to current paradigms, implemented in engineered systems such as spiking neural network hardware architectures. We do not expect that DN, as a computational module, is necessarily topmost in the computational hierarchy and therefore we expect behavioral correlates to potentially be indirect. More emphasis is placed here on discussing DN as a fundamental problem-solving neural machine as opposed to relating its solutions to possibly convoluted human behavioral patterns.

## 2 Results

### 2.1 Building blocks of Darwinian neurodynamics

When proposing a DN architecture, one has to specify how necessary elements of a Darwinian evolutionary process, namely, replication and selection of units of evolution, and heritable variation over them, are neurally implemented. In the following, we show how these choices can be made by building up our model architecture, based on dynamic firing patterns as evolutionary units, which we refer to as *recurrent DN*, in a step-by-step manner.

#### Unit of evolution

The unit of evolution in recurrent DN is the output firing pattern of a recurrent computational unit, as illustrated in Figure 1a. It is, in the current model, a one-dimensional time-dependent signal, usually interpreted as neuronal firing rate. Importantly, the unit of evolution is not the recurrent network itself, it is the computation it does. We use reservoir computing units to implement these recurrent computations. In reservoir computing units, learning takes place only through the linear readout synapses; the recurrent connections within the network are fixed. This makes the map from computer to computation highly degenerate: the same computation, i.e., output signal, can be represented (at least approximately) by an enormous number of different reservoirs. The reservoirs themselves form representational slots for the replicators, i.e., the output signals.

**Figure 1:**
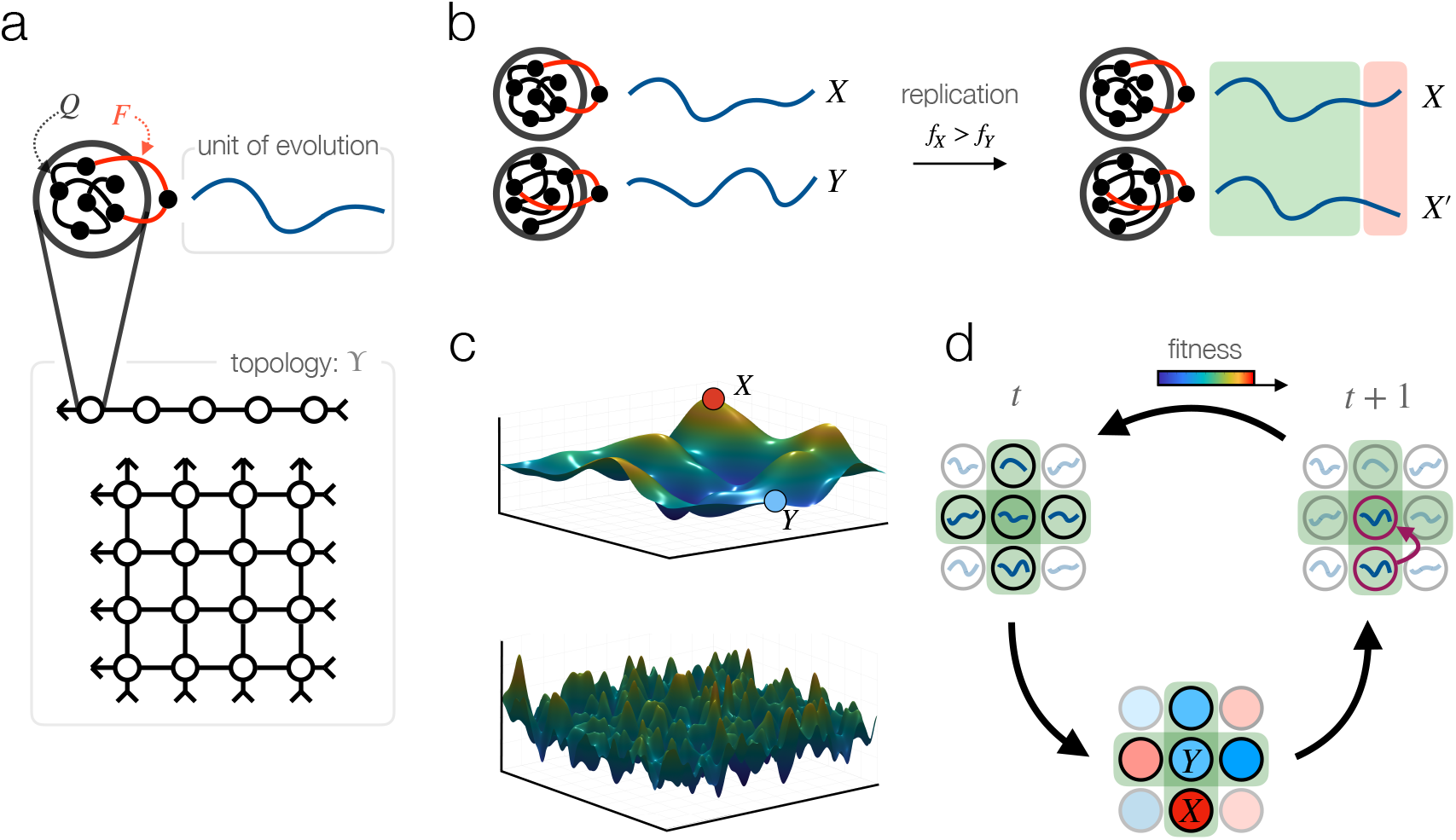
Schematic illustration of component processes of recurrent Darwinian neurodynamics. a. Reservoir computers, arranged in a 1D array or a 2D lattice, host replicating signals. b. Replication of output signals takes place by one reservoir training another. Reservoirs that output fitter signals train, those that output less fit signals learn. Training time and neural noise determine the fidelity of replication. c. Illustration of ruggedness of fitness landscapes. More rugged landscapes, such as the one in the bottom, has more non-equivalent local maxima, making optimization more difficult. d) Full dynamics. During one generation, each reservoir output *Y* is compared to those of its neighbours; the reservoir with the best output signal among the neighbours (here, *X*) replicates its signal to reservoir *Y* by training it.

#### Replication

The crucial step in any DN architecture is the replication of firing patterns, utilizing local learning rules. Here, in recurrent DN, replication takes place by one recurrent network training another, as shown in Figure 1B. More precisely, the training signal of reservoir A is the output of reservoir B, rendering, after some training period, the two output signals *approximately* the same. In other words, the output of reservoir B serves as a template for the output of reservoir A, resulting in the replication of the output of reservoir B, with some variation. We use the FORCE algorithm to train reservoirs. Note that the FORCE algorithm is a supervised learning algorithm, which is in general inapplicable for computations where a supervised signal is not available; an evolutionary process artificially creates supervised signals by displacing the low-dimensional informational bottleneck between the environment and the system from reservoir training to fitness computations.

#### Heritable variation

Heritable variation is a necessary for selection to act upon, and therefore, to improve solutions over time according to their fitness metric. Variation comes from two sources in our model. *i*) It comes from imperfect training of one network by another via the FORCE algorithm, and *ii*) it also comes from neural noise that we model as white noise added to the learned signal. Importantly, this variation is heritable if the resulting mutated signal can be learned by yet another reservoir. If the set of heritable signals is practically unlimited (i.e., much larger than the population size, the number of replicating signals), the system exhibits unlimited hereditary potential, upon which long-term evolution can take place.

#### Fitness landscape

Evolution, like reinforcement learning, necessarily channels all information about the environment through a one-dimensional bottleneck. It is this scalar-valued function, the fitness function, that must contain all task-related information if the architecture itself is meant to be of general purpose. It is not in the scope of this paper to discuss possible implementations of this evaluation signal in wet brains, but this similarity suggests that if Darwinian dynamics takes place in brains, the striatal reward system might be a possible candidate location of fitness assigment. In this paper, we intend to demonstrate the feasibility of an emergent Darwinian process over dynamic neural firing patterns in an as task-agnostic manner as possible. We therefore evaluate our recurrent DN architecture on general combinatorial landscapes with many non-equivalent local optima: on the Tavelling Salesman Problem and on NK-landscapes with different ruggedness. Figure 1c visualizes the idea of ruggedness of a continuous 2-dimensional landscape; in our simulations, we use high-dimensional discrete landscapes that are difficult to visualize.

NK-landscapes [33], originally modelling epistatic fitness interactions of genes, are simple non-deterministic landscapes that are able to model arbitrary level of ruggedness, set by a parameter *K*. They assign fitness *f* to binary sequences *x*_1_, *x*_2_, … *x_N_* as

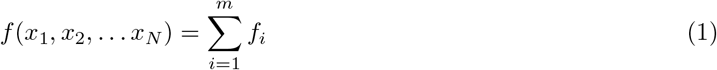

where each *f_i_* fitness-component depends on exactly *K* ≤ *N* coordinates. The value of these fitness components is drawn from the uniform distribution over [0,1] independently. As an effect of the stochastic nature of NK landscape generation, realizations of landscapes with the same parameters *N* and *K* generally differ. *N* and *K* therefore define task *classes*, with *N*, the number of bits or “genes”, parametrizing problem size and *K* parametrizing “ruggedness” or “frustration” in the system. Note that NK landscapes include spin glass Hamiltonians as special case, corresponding to (a subset of) *K* = 2 landscapes. *K* > 2 landscapes, on the other hand, model higher order interactions between elements. Overall, NK landscapes form a simple *and* flexible benchmark for combinatorial problems.

#### Selection

Selection in DN refers to the assignment of number of offspring to replicating firing rate patterns. It is done through evaluating the solutions represented by the replicating signals according to a scalar function, the fitness. In our architecture, reservoirs that output higher-fitness signals become *trainer* reservoirs, reservoirs outputting lower-fitness signals are assigned to be *learners*. The signal of trainer reservoirs reproduce, while the signal of learner reservoirs dies (i.e., disappears form the population of signals). The full evolutionary process over a two-dimensional sheet of reservoirs is illustrated on Figure 1d.

#### Population structure defined by cortical topology

Fitness values determine the reproduction rate of signals, in terms of assigning the corresponding recurrent unit to be either teacher or learner. This assignment is dependent on comparing a subset of signals with each other. We define these competing subsets in terms of a connection topology of recurrent units: each recurrent unit (which is a neural network itself) is competing with its direct neighbours according to this pre-defined meta-topology. Special cases of this network meta-topology include a one-dimensional array of networks for illustration purposes (see phylogenetic trees of signals in section 2.2), and a two-dimensional grid, that might be identified with a cortical sheet of canonical microcircuits, see Figure 1a. Although these meta-topologies induce a local competition dynamics, competing groups overlap and therefore high-fitness solutions are able to spread in the population globally. Population structure, defined here by this meta-topology of recurrent units, strongly shapes the course of evolutionary competition dynamics in general. If the evolutionary system as a whole is itself subject to optimization, e.g. by another evolutionary process at a higher level, we might expect to observe fine-tuned hyper-parameters in general, and a fine-tuned cortical topology in particular. Although optimizing cortical topology for various performance measures is out of the scope of this paper, we refer to evolutionary graph theory [34] as a systematic study of amplifying and diminishing effects of population structure on selection.

#### Representation

Notice that replication acts at the level of signals whereas selection acts at the level of solutions represented by the signals. This mapping between signal to solution determines the accessibility of solutions by evolution, just like the genotype-phenotype map in biology determines the accessibility of phenotypes. Solutions that are represented by many signals can be found with relative ease by evolutionary trajectories, whereas solutions that are represented by a tiny fraction of signal space or not at all are difficult to find. Statistical properties of the signal-solution map are therefore of fundamental importance. Since our goal here is to explore how well our recurrent DN arcitecture performs over rugged fitness surfaces with a large number of non-equivalent local optima, we map the signals to combinatorial solutions represented by *i*) permutations, in case of the travelling salesman problem (TSP) fitness landscape, or by *ii*) binary sequences, in case of the NK-landscape. These signal-solution maps are visualized in Figure 2a; their statistical properties are shown in Figures 2b-2e.

**Figure 2:**
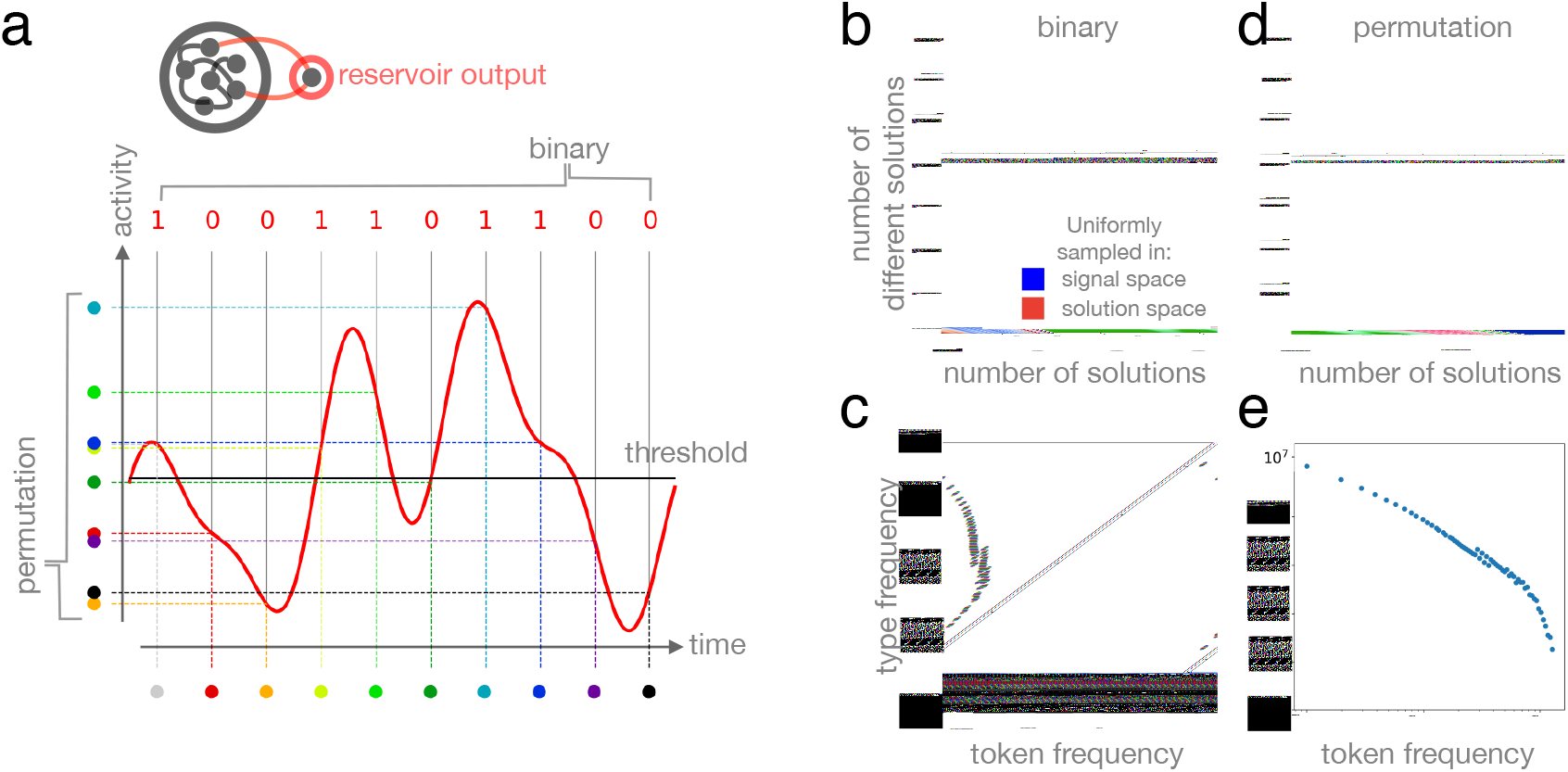
Signal to solution maps and their statistical properties. a. Output signals of reservoirs, shown in red, are mapped to i) binary sequences by thresholding signal amplitude (i.e., neural activity) at regular time intervals (top), or to permutations by ordering these signal amplitudes. b,c,d,e. Statistical representation bias of signal to solution maps, showing that a few solutions are represented by many signals, while most solutions are represented by only a few signals. b, d. Number of different solutions as a function of the number of sampled solutions. Samples are drawn uniformly in signal space (blue; see Methods for details), or in solution space (red). While uniform sampling in solution space saturates exponentially, uniform sampling in signal space leads to a different saturation curve as an effect of some solutions being sampled many times while others are sampled less or not at all. c,e. Number of different solutions (types, *y* axis) with a given sampled frequency (token frequency, *x* axis), mirroring representation bias.

### 2.2 Putting the parts together: evolution in one or two cortical dimensions

Having described the essential ingredients, we now focus on exploring the characteristics of recurrent Darwinian neurodynamics as a coherent computational framework. In all cases, we either use a one dimensional array, or a two-dimensional sheet of reservoirs as meta-topology (i.e.., population structure from an evolutionary point of view), mirroring the possibility of representation by cortical microcircuits. First, we illustrate the evolution of the *best* firing pattern and the corresponding solution on a well-known combinatorial optimization problem, the traveling salesman problem (TSP), see Figure 3a. We then turn to a more general (but less visual) set of combinatorial problems, optimization over NK landscapes. We introduce an analysis technique for tracking the evolution of firing patterns as a population, *neural phylogenies* (Figure 3b, 3c), and demonstate the ability of recurrent DN to continuously improve upon currently best solutions through beneficial mutations (Figure 3d). Next, we formulate fitness in terms of *information gain* in order to make evolution over different landscapes comparable, and we show the time evolution of fitness distribution of the population over different landscapes and initial populations (Figure 3e). Finally, we demonstrate the existence of a sharp neural-evolutionary *error threshold*, a neural noise level above which information gained by evolution cannot be maintained (Figure 4a,b,c).

**Figure 3:**
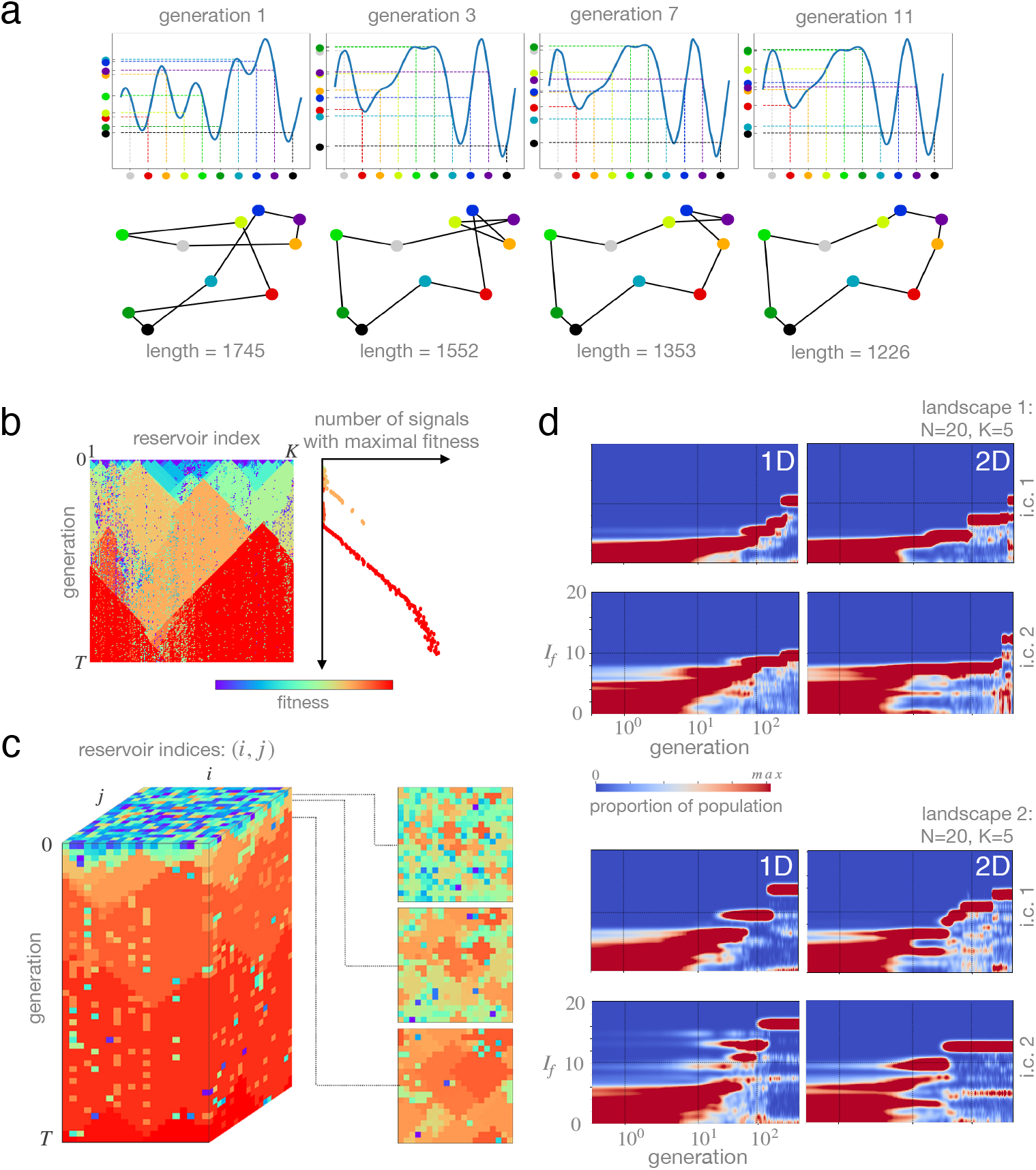
a. Recurrent Darwinian neurodynamics over a 1D array of reservoirs solving the travelling salesman problem (TSP). Shorter paths in solutions space correspond to higher fitness in signal space. Top: Highest-fitness signals of their generation. Bottom: Corresponding solutions, permutations representing a path over major Hungarian cities. b. Phylogenies of neural firing patterns. A 1D array of reservoirs is shown horizontally; time evolution proceeds vertically downwards. Color corresponds to the fitness of signals outputted by their reservoirs, evaluated over an NK landscape with *N* = 20 and *K* = 5. Signals with higher fitness spread over local connections between reservoirs. Reservoirs differ in their ability to learn, in extreme cases, they form defects or local walls. If such a mechanism operates in wet brains, such defectuous units might have been under strong selection pressure over timescales of biological evolution. c. Phylogeny of firing patterns over a 2D sheet of reservoirs. Cross sections (right) illustrate local spread of high fitness firing patterns, leading to competing homogeneous islands. Sporadic reservoirs with diminished ability to learn do not hinder the process significantly, as opposed to the 1D case. In both 1D and 2D cases, learner reservoirs inherit the signal of the teacher reservoir with some variation. The level of variation, i.e., neural mutation rate, is set by i) training time and ii) neural noise. d. Evolutionary information gain *I_f_* over time. *I_f_*, a comparable measure of fitness over different landscapes, corresponds to the 2*^−I_f_^* th highest fitness value in the landscape. In all cases, the population of firing patterns adapts to an NK landscape with *N* = 20 (hence the maximum of *I_f_* is 20) and *K* = 5, implying a high level of ruggedness and correspondingly, low information content of local gradients. We display three levels of between-run variability here. i) dimensionality of the arrangement of reservoirs (columns); ii) two realizations of an NK landscape with *N* = 20 and *K* = 5 (top & bottom); iii) two different initial populations of firing patterns (rows in each block). Although there is a high level of between-run variability across all dimensions, the population of firing patterns keeps finding better and better solutions in all cases, suggesting the feasibility of this process as an efficient computational module.

**Figure 4:**
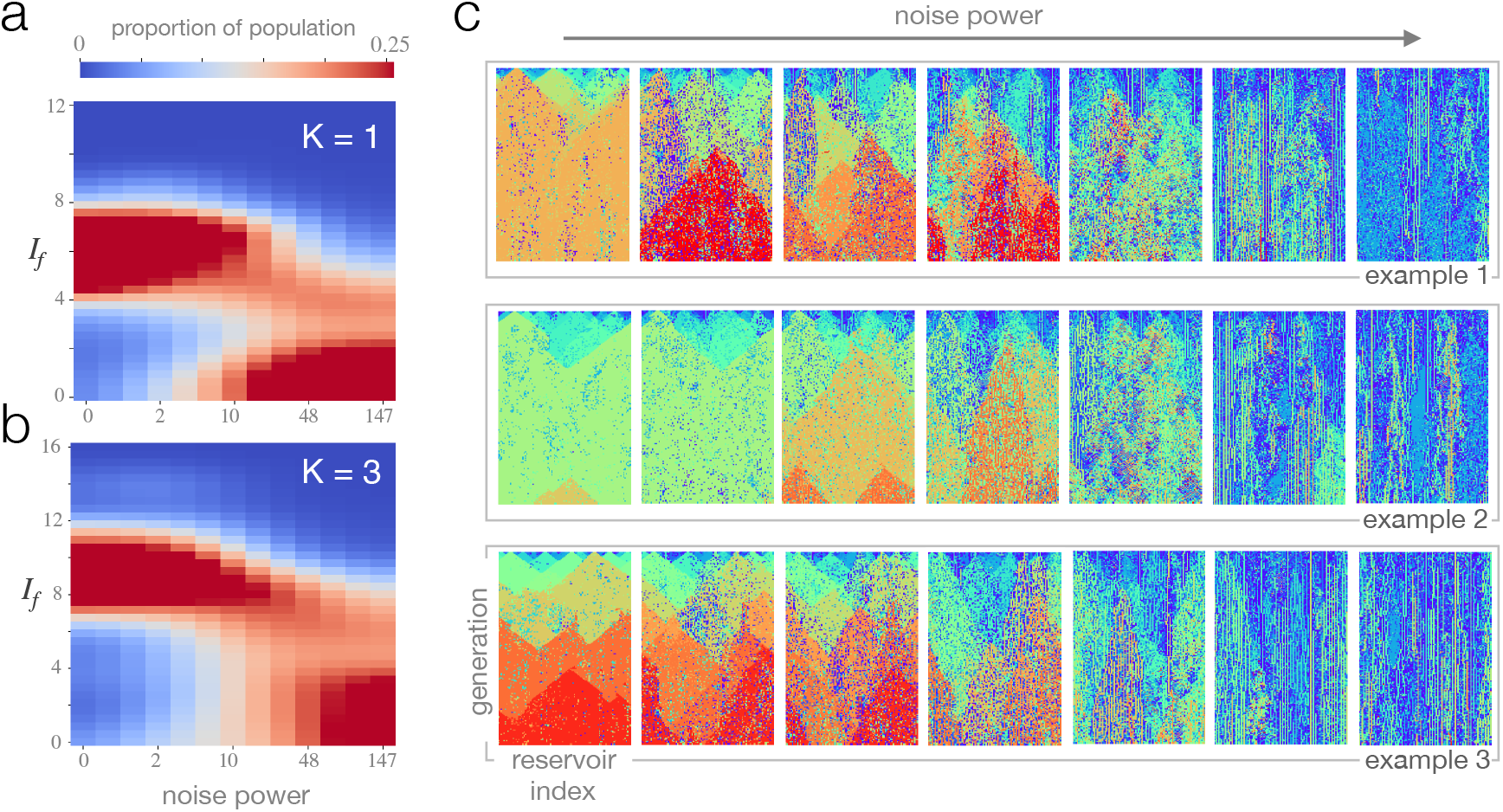
Error threshold for Darwinian neurodynamics, a critical neural noise level beyond which information accumulated by the evolutionary process cannot be maintained. We posit that this threshold defines the regime (in any Darwinian neurodynamics architecture) where a replicator-based understanding of neural computations might provide significant insight. a,b. Information gain distribution *I_f_* of the 200th generation, in the function of increasing neural noise. Each column (corresponding to a given neural noise level) is averaged over 5 runs. In both *K* = 1 (a) and *K* = 3 (b), information content accumulated by evolution suddenly drops to a practically random level. The critical noise value at which this transition happens defines the error threshold for Darwinian neurodynamics. c. Neural phylogenies as more and more noise is added. All runs are evaluated over the same NK landscape with *N* = 20 and *K* = 3; rows differ in their initial population of signals. Within rows, only the level of added noise varies. Above the error threshold, high fitness signals disappear from the population before they could undergo beneficial mutations.

#### Breeding Traveling Salesmen

Given a list of cities, the Traveling Salesman Problem (TSP), asks for the shortest length route that visits each city and returns to the origin. While simple to state, both the rapid growth of the number of possible routes (permutations of the city list) with the number of cities and the difficulty of factoring the problem into simpler sub-problems renders the TSP NP hard. A wide range of algorithmic approaches have been proposed for the TSP, including those inspired by evolutionary dynamics. Here, we select the TSP as a classic example of a difficult combinatorial optimization problem to demonstrate the behaviour of this recurrent DN architecture. The goal of this illustration is not to compete with other specialized algorithms employing TSP specific heuristics, but to highlight the importance of representation and demonstrate the capacity of this DN system.

Figure 2a illustrates one possible encoding of a permutation in a time varying signal; here each coloured dot represents a city identity and the order along the *y* axis represents a route. Notice again that many other possible signals (genotypes) are consistent with a specific route (phenotype) - this degeneracy may have important consequences for the *evolvability* of the representation (see Discussion). Figure 3a shows a toy example TSP with 10 cities (a solution space size of ~ 2^18^) that is solved via evolutionary dynamics over a 1D array of reservoirs.

Although performance, in general, depends on the initial signals and the inner weights of reservoirs (determining the scope of signals that can be learned by each reservoir), the single run we visualize in Figure 3a captures an essential feature of any evolutionary dynamics: the automatic reallocation of computational resources to improving upon currently best solutions. After pooling a diverse set of initial conditions, the dynamics automatically narrows down to local improvements around highly promising solutions, culminating, in this example, in the global optimum after 11 generations.

#### Phylogeny of firing patterns

Although tracking the best solution at all times already offers intuition behind our neuro-evolutionary dynamics, a higher resolution analysis can be given by following the fate of *all* signals as they replicate and mutate, leading to neural *phylogenies* (ancestry trees), visualized in Figures 3b and 3c.

We twist conventional phylogenies in two ways. First, since the spatial arrangement of reservoirs (i.e., hosts of replicating signals) is fixed to be a 1D array or a 2D lattice, we simply use this additional information to fix the spatial position of replicators at all times to be those of their host reservoirs. As time (measured in the number of generations) passes vertically, a vertical line corresponds to replicators in the same host reservoir at subsequent generations. Second, in order to keep phylogenies simple, we visualize only the *fitness* of each signal at all times; since the fitness of different signals is likely to be different, this provides a good proxy for the actual phylogeny of signals.

Such neural phylogenies are especially useful in detecting the dynamics of relevant scales of competition. Initially, out of the sea of randomly chosen initial conditions, locally good signals emerge and start to spread. They then form growing homogeneous islands. Higher-fitness islands out compete lower-fitness ones, the relevant scale of competition increases. In the meantime, computations given by local fitness comparisons are allocated to compare higher and higher fitness solutions. Phylogenies also reveal the reservoirs’ differential ability to learn; in extreme cases, particularly inflexible reservoirs form a local wall, preventing otherwise high-fitness signals as replicators to spread.

#### Information gain

Since fitness values *f* cannot be compared across landscapes, we transform fitness to *information gain I_f_*, calculated as *I_f_* = – log_2_ *q_f_*, where *q_f_* is the fraction of all types (here, binary sequences of length *N*) that have higher or equal fitness than *f*. If the fitness *f* of the current type is the median of all fitness values, then *I_f_* = 1 bit; if *f* is at 75 percentile, *I_f_* = 2 bits, and so on. Figure 3d shows the distribution of information gain *I_f_* in the population as it evolves over NK landscapes with *N* = 20 and *K* = 5. We compare runs over three axes: different population of initial signals, different landscapes, and the dimensionality of the meta-topology of reservoirs (i.e., 1D array or 2D lattice). The high observed between-run variability makes it uninformative to average performance over multiple runs, we therefore present results of single runs. In all runs, information gain increases monotonously. It increases in jumps, corresponding to rare beneficial mutations that improve the currently best solutions. Between jumps, information is maintained: highest-fitness solutions do not disappear from the population unless a higher-fitness mutant appears. This is marked by relatively steady periods of information gain distributions. Finally, islands of lower-fitness solutions are sometimes also kept by the dynamics, as the existence of multiple relative stable modes of the information gain distribution suggests.

#### Error threshold for Darwinian neurodynamics

The *error threshold* in population genetics refers to a critical mutation rate above which populations diffuse out of fitness peaks to neighbouring genotypes [35]. Although these neighbouring genotypes have lower fitnesses, they are more numerous; lower mutation rate allows for i) narrower or ii) lower fitness peaks. Beyond the error threshold, locally good types are not maintained, and consequently, cannot be improved upon by beneficial mutations.

In analogy with population genetics, here we present an error threshold for Darwinian neurodynamics: a critical neural noise value above which the Darwinian evolutionary process over neural replicators breaks down. As shown in Figures 4a and 4b, there is indeed a sharp transition from high to low fitness, and correspondingly, high to low information gain *I_f_* as neural the noise level increases.

This threshold might provide a useful definition to what *neural replication* means: an approximate copying of firing patterns on top of which a functional Darwinian process, below error threshold, can be built. For such processes, theoretical frameworks and arguments from evolutionary biology and population genetics might be efficiently transferred to understand neural dynamics on a computational and algorithmic level.

## 3 Discussion

Darwinian neurodynamics links two well-studied and experimentally rooted conceptual systems: neuroscience and evolution, with the increasingly powerful models and tools of machine learning. The endeavor has two main goals: i) to answer ascertain whether or not brains use principles of Darwinian evolution to select actions or representations, and if so, to understand how these principles they realized in neural populations; ii) to assemble a conceptual bridge over which evolutionary principles can be transferred to neuromorphic learning architectures. These two goals might very well overlap. This paper approaches goal i) from a reverse engineering perspective, asking how local learning in populations of neurons can sustain a higher level evolutionary dynamics. We discuss the key theoretical issues involved in mapping Darwinian dynamics onto neuronal populations, anchoring our discussion in a simple computational paradigm based on the emergent replication of dynamic firing patterns and pointing at the series of decisions one has to make when setting up a DN architecture all the way down to algorithmic and implementational choices.

The main conceptual issues, illustrated by the recurrent DN architecture introduced in this paper, are the following. Natural selection is an algorithm that is capable of dynamical reallocation of computational resources from globally bad to globally good solutions. It does it in a simple and therefore universally robust way: it copies good solutions in place of bad ones. Many engineering methods are inspired by this idea, notably evolution strategies and estimation of distribution algorithms [36, 37] - both iteratively propose new solutions based on the goodness of previously proposed ones. These, as all ML algorithms, trade off universality for efficiency in particular cases [38]. Darwinian neurodynamics is therefore a robust, generalist solution to the dynamic reallocation problem. Evolution as a dynamic reallocation algorithm, however, is only useful if existing solutions are not washed away by the sea of mutations while searching for even better ones. In other words, although mutations generate the necessary variation upon which selection can act, there is an upper limit, given by the Eigen error threshold in simple combinatorial systems. Mutations generate variation through which the space of possible solutions is explored. If replication and selection act at different levels, connected by e.g. a genotype-phenotype or, in our case, a signal-solution map, then generated variation at the replication level is transformed in the selection level. This leads to a potentially highly non-uniform exploration of phenotypes/solutions. These non-uniformities can take two forms. *i*) Evolvability/facilitated variation: generated variation at the phenotype level is informed by the evolutionary past and proposes solutions that have a higher expected fitness than proposed solutions without this mechanism. *ii*) Limited accessibility of phenotpic regions: phenotypes cannot be represented with equal ease. This effect combines with fitness to determine evolutionary trajectories.

Furthermore, going down in the algorithmic hierarchy to the neural representation of Darwinian dynamics, the recurrent DN model we present in this paper exemplifies another core idea: the use of a population of *supervised* learning modules to solve *reinforcement* learning like problems, where information from the environment is only available through a scalar bottleneck, called reward or fitness. Our architecture minimally couples these two dynamics: the role of being a teacher or learner unit is determined via a simple comparison of reward/fitness; variation on which the Darwinian process feeds is introduced ‘blindly’ through the necessarily imperfect supervised training. There is no need for gradient information or another computational module that is aware of the problem structure; the population of agents - in this case, neural firing patterns - explores both locally and globally via emergent Darwinian dynamics. We envisage that the architecture we present here can be substantially generalized and extended along the lines of these core ideas to more realistic scenarios. In particular, copying (noisy) higher-dimensional attractors instead of one-dimensional time-varying signals is one such possibility.

One might expect that brains, having evolved under tight energy and space constraints, would employ evolutionary processes, including the maintenance of a population of solutions, only if it provides a qualitative advantage in search problems over non-population based processes. What are these qualitative advantages? Here we provide a subjective account, emphasizing algorithmic features that are unique to evolution, and pointing at their relation to cognitive problem solving.

A key feature of evolution is the unrestrictedness of representations, and the opportunity for open-endedness this unrestrictedness offers [39, 40]. Since there is no need for estimating how to improve on the current representation, e.g. by computing gradients or solving complicated credit assignment problems, in evolution, “anything goes”. Anything that is, from which assembly instructions can be copied. The more the phenotypes (or candidate solutions, in the language of DN) are decoupled from the replicating substrate that holds the set of instructions for making the phenotype, the more freely the (infinite) phenotype space can be explored. In biology, phenotypes are constructed from genotypes by (probabilistic) developmental programs; in this sense the phenotype is decoupled from the genotype. These programs are sets of instructions that encode incremental and modular construction, they allow for arbitrary re-use and can modify existing modules, paving the royal road to open-endedness. This royal road may well have been stumbled upon by neural systems at some point in their convoluted evolutionary history, replete with complex problem solving. Indeed, probabilistic programs leading to compositional/hierarchical action sequences and the inference of these programs from sensory data offers a successful modeling framework for human cognition [41, 42].

Biological evolution goes beyond this, however, by employing an array of clever tricks to boost search in (developmental) program space. Tricks that are not tied to the specificities of life as we know it. These include i) a variety of methods for maintaining a diverse set of solutions [43, 44], ii) (re)combining already invented modules at different timescales [45, 46, 47], and iii) accelerating change by setting up arms races of various kinds [48, 49]. We believe that exploring these substrate independent algorithmic tricks in the context of Darwinian neurodynamics potentially holds unexpected insights along the way.

## 4 Materials and Methods

### 4.1 Reservoir computing model

#### Unit of evolution

In this model of DN, the unit of evolution is the output activity of a reservoir computer. A reservoir computer, *R* consists of i) a reservoir containing *n_reservoir_* point neurons (*n_reservoir_* is typically chosen to be relatively large, such that inputs can be represented in a high dimensional space of activities), and ii) a readout consisting of *n_readout_* neurons. The recurrent weights between units in the reservoir are contained in matrix ***Q*** and the feedforward weights from the reservoir to the readout in matrix ***F*** (see Figure 1a, where *n_reservoir_* = 6 and *n_readout_* = 1). The readout layer is fed back to the reservoir via feedback weights contained in matrix ***B***. Only the readout weights, ***F*** are trained; the recurrent and feedback weights remain fixed throughout training [50].

***Q*** is a sparse matrix, with connection probability *p*, and zero entries along its diagonal. Weight matrices are initialized as follows:

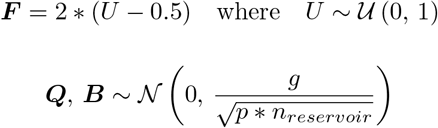

where *g* is a scaling constant that controls the presence and/or strength of spontaneous chaotic activity in the network.

The output of reservoir *R_i_* is signal ***y**^i^* - these signals are the units of evolution (the genotypes) and are parameterized by their respective reservoirs. In the examples considered here, ***y**^i^* is a 1D vector.

In general, reservoir dynamics are described by the following differential equation.

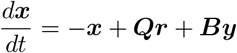

where ***x*** is the *n_reservoir_*-dimensional vector of neuronal activities within the reservoir, ***r*** = tanh(***x***) is the vector of neuronal firing rates, and, again, ***y*** = ***Fr*** is the output of the reservoir (the activity of the readout neuron).

A number, *N_res_*, of reservoir computers can be connected to form a network in which the output of one reservoir may act as the input of another. Each node in this network is therefore a reservoir computing unit, *R_i_* where *i* = 1,…, *N_res_*, and an edge between nodes *i* and *j* represents the idea that *R_i_* and *R_j_* may each either receive input from or send output activity to the other. In this sense one network may act as a *teacher* and the other a *learner*. Note that we describe a network of networks, where the higher level connectivity between units *R_i_* is denoted 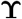 and the lower level connectivity describes the recurrent, output and feedback weights (contained in matrices ***Q**^i^, **F**^i^* and ***B**^i^* respectively) in each *R_i_* For 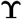, we consider one and two dimensional regular lattices where only neighbouring reservoir units are connected - in the 1D case, each node is connected to two others and in the 2D case, to four others (the set of neighbouring nodes to a given node *a* is denoted *X_a_*). These lattices wrap around such that there are no boundaries.

#### Initialization

In a typical run, each reservoir unit *R_i_* is initialized by training the output weights ***F**^i^* such that *R_i_* outputs a randomly assigned signal. This initialization signal, ***y***^*i*, 0^, is constructed by drawing *n_Fourier_* sine and cosine coefficients from a uniform distribution in the interval [–0.5,0.5]. The FORCE algorithm is used to train output weights *F_i_* and is described in [50].

#### Replication

In the same manner, in any given generation *t*, each *R_i_* can act as a *learner*, i.e. learn the output of another unit *R_j, j≠i, j∈X_i__*, or as a *teacher*, where output weights of the learner unit are trained via FORCE such that the output activity pattern is copied from teacher to learner (***v**^j,t^* → ***v***^*i,t*+1^).

FORCE algorithm is a supervised learning method, therefore it is necessary to provide an explicit target signal. Here the output signal ***y**^j^* acts as the target for *R_i_* during training. Using the recursive least squares (RLS) algorithm, the learning rule is as follows

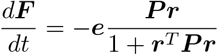

where the error vector, ***e***, is

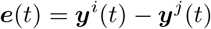

and ***P*** is an *n_neurons_ × n_neurons_* matrix which is the running estimate of the inverse of the correlation matrix of firing rates plus a regularization term. It is updated by the following rule

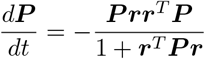

The initial value of ***P*** is

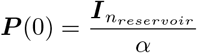

where ***I**_nreservoir_* is the identity matrix and *α* acts as the learning rate of the FORCE algorithm.

#### Selection

In this model, the direction in which output patterns are copied is determined by the relative fitness of the two signals. Each signal-genotype is mapped to a solution-phenotype via a neural genotype-phenotype (GP) map. Here we consider two deterministic GP maps: i) a signal vector, ***y**^i^* is mapped to a binary string, ***v**^i^*, of length *N*, where *i* indexes the signals (see Figure 2a above), and ii) ***y**^i^* to permutation vector ***h**^i^* (see Figure 2a along the activity axis), where the ordering is defined by the magnitude of the signal at bin edges. The fitness landscape is a map from the vector valued phenotype, either ***v**^i^* or ***h**^i^* to a scalar fitness, *f^i^*. This mapping depends on the problem under consideration - the TSP and NK models are two examples described in this paper.

#### Variation

The copying of signals from one reservoir to another is typically imperfect - this process introduces ‘mutations’ into the copied signal, providing variation upon which selection can act. The quality of copying between reservoir units varies significantly between signals; unfortunately, this is not readily controlled with the FORCE algorithm. One approach to controlling the mutation rate involves injecting additional noise into the copied signal (see Figure 4). Here, *white* noise of varying strengths is added to the signal of the learner after each copying event.

#### Darwinian dynamics

A generation is defined as one round of signal copying; in this round each reservoir output signal is compared with its neighbouring reservoirs (the indices are members of *X_i_*). In the 1D examples (see Figure 3b), 2*N_res_* comparisons are made in parallel each generation as reservoir *R_i_* is compared to *R*_*i*−1_ and *R*_*i*+1_. In the 2D networks from Figure 3c, this number is 4*N_res_*, as each reservoir is compared to its neighbours on the square grid, 2-2 by each dimension. In the case of multiple reservoirs with maximal fitness in the neighbourhood of *R_i_*, the teacher for *R_i_* is selected at random from those with maximal fitness. The network is run for a total of *T* generations.

### 4.2 Fitness landscapes

#### Traveling Salesman Problem

The traveling salesman problem can be simply stated as: compute the shortest possible route that visits each city (from a given list of destinations) and returns to the origin city. In this classic NP hard combinatorial optimization problem, the total path length (*l_i_*) for a solution vector ***h**^i^*, can be related to the scalar fitness, *f_i_* by a function such as *f_i_* = – *l_i_* or *f_i_* = 1/*l_i_*.

#### NK landscape models

In the NK models, the mapping of an input (here a binary string ***v***) to a scalar fitness can be tuned to make the landscape more or less difficult to traverse via local gradients. The *N* from NK is the length of the binary sequence that is evaluated; the size of the space of binaries is therefore *M* = 2*^N^*. *K* on the other hand controls the extent of epistasis (interaction between positions of the binary string) and therefore the ruggedness of the fitness landscapes [51]. Hard search problems are characterised by very large search spaces and complex fitness landscapes with many local optima. Even low levels of epistasis (*K* > 1) generally makes finding local optima within the high dimensional maze difficult [52].

### 4.3 Simulation parameters

Those parameters that were fixed throughout all simulations shown in the paper are described in Table 1, while the parameters that were varied between simulations are described in Table 2 below.

**Table 2:** Fixed parameters

|  |  |  |
| --- | --- | --- |
| Number of neurons | $n_{reservoir}$ | 1000 |
| Number of readout neurons | $n_{readout}$ | 1 |
| Connection probability of sparse $\mathbf{Q}$ matrix | $p$ | 0.1 |
| Chaocity parameter | $g$ | 1.5 |
| Timestep | $dt$ | 0.1 |
| Signal generation time | $t_{signal}$ | 300 |
| Learning rate of FORCE algorithm | $\alpha$ | 1 |
| Training time | $t_{train}$ | 300 |
| Evaluation time | $t_{evaluation}$ | 30 |
| Number of sine and cosine coefficients of initial signal | $n_{Fourier}$ | 5 |

**Table 3:**
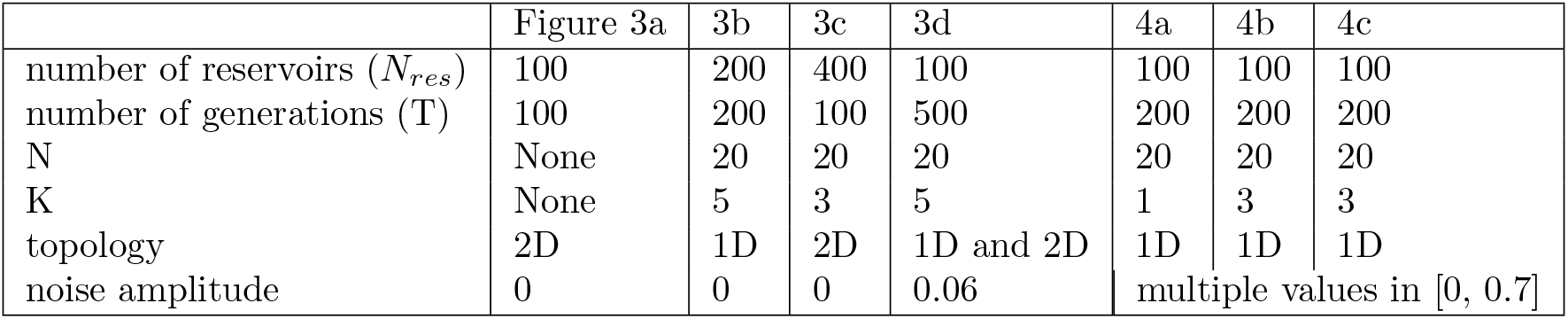
Parameters that are varied

|  | Figure 3a | 3b | 3c | 3d | 4a | 4b | 4c |
| --- | --- | --- | --- | --- | --- | --- | --- |
| number of reservoirs ($N_{res}$) | 100 | 200 | 400 | 100 | 100 | 100 | 100 |
| number of generations (T) | 100 | 200 | 100 | 500 | 200 | 200 | 200 |
| N | None | 20 | 20 | 20 | 20 | 20 | 20 |
| K | None | 5 | 3 | 5 | 1 | 3 | 3 |
| topology | 2D | 1D | 2D | 1D and 2D | 1D | 1D | 1D |
| noise amplitude | 0 | 0 | 0 | 0.06 | multiple values in [0, 0.7] |  |  |

Link to the open source Python code: https://github.com/csillagm/reservoir-dn

